# PITA, a pre-bilaterian p75NTR, is the evolutionary ancestor of TNF receptors

**DOI:** 10.1101/2021.12.26.474206

**Authors:** Mark J. Cumming, Julien Gibon, Wayne S. Sossin, Philip A. Barker

**Author notes:** Corresponding author, (PAB).

## Abstract

Tumor necrosis factor receptors (TNFRs) regulate a diverse array of biological functions, including adaptive immunity, neurodevelopment, and many others. Although TNFRs are expressed in all metazoan phyla, a coherent model of the molecular origins of mammalian TNFRs—and how they relate to TNFRs in other phyla—has remained elusive. To address this, we executed a large-scale, systematic Basic Local Alignment Search Tool (BLAST)-based approach to trace the evolutionary ancestry of all 29 human TNFRs. We discovered that all human TNFRs are descendants of a single pre-bilaterian TNFR with strong sequence similarity to the p75 neurotrophin receptor (p75NTR), which we designate as PITA for ‘*p75NTR is the TNFR Ancestor*’. A distinct subset of human TNFRs—including EDAR, XEDAR and TROY—share a unique history as descendants of *EDAR-XEDAR-TROY* (EXT), which diverged from PITA in a bilaterian ancestor. Most PITA descendants possess a death domain (DD) within their intracellular domain (ICD) but EXTs do not. PITA descendants are expressed in all bilaterian phyla and Cnidaria, but not in non-planulozoan ParaHoxozoa, suggesting that PITA originated in an ancestral planulozoan. *Drosophila melanogaster* TNFRs (Wengen (Wgn) and Grindelwald (Grnd)) were identified as divergent PITA descendants, providing the first evolutionary link between this model TNFR system and the mammalian TNFR superfamily. This study reveals PITA as the ancestor to human and *Drosophila* TNFR systems and describes an evolutionary model that will facilitate deciphering TNF-TNFR functions in health and disease.

## Introduction

In 1893, immunologist Dr. William B. Coley induced stable remission of an inoperable sarcoma by injection of a bacterial immunogen (‘Coley toxin’) directly into the primary tumor (1). This anti-tumoral immune response was rediscovered 69 years later, with the discovery that sera derived from mice exposed to a bacterial polysaccharide could induce tumor regression *in vivo*, suggesting that a ‘tumor necrotizing factor’ must be present in the serum (2). This factor was subsequently isolated and named tumour necrosis factor (TNF; now TNFα) (3). Over the past decades, 19 human TNF ligands and 29 TNF receptors (TNFRs) have been characterized in humans [reviewed in (4)]. TNFRs mediate a diverse array of biological functions and are now known to function as critical regulators of innate immunity (5,6); adaptive immunity (7,8); and nervous system development (9–15) and maintenance (16–18). Aberrant TNFR signaling is implicated in viral infections (19), autoimmune disorders (20,21), neuropsychiatric disorders (22–24), neurodegenerative disease (25–31), and other disease states (32,33).

TNFRs are non-enzymatic single-pass transmembrane proteins that signal via ligand-dependent and -independent recruitment of adaptor proteins to their intracellular domain (ICD), most notably the TNFR-associated factor (TRAF) family of ubiquitin ligases (34). TNFRs are structurally related by the presence of one or more cysteine-rich domains (CRDs) in their extracellular domain (ECD). The CRD is ~40 amino acids (AA) in length and contains 6 conserved cysteine residues, which form a specific pattern of disulfide bridging (C_1_-C_2_, C_3_-C_5_, C_4_-C_6_) that provide structural rigidity to the domain. Extracellular CRDs are typically arranged in tandem arrays that produce distinct ‘CRD clusters’—with an overall primary sequence unique to each TNFR—that mediate ligand binding. Most TNFRs exist as homotrimers (35) and signal in a 3:3 TNF:TNFR stoichiometry, with the exception of p75NTR. p75NTR does not bind a TNF ligand, but instead binds neurotrophin (NT) ligands (NGF, BDNF, NT-3 and NT-4/5) in a 2:2 NT:p75NTR stoichiometry (36). p75NTR also binds the pro-form of NGF if the Vps10p family member sortilin is present as a co-receptor (37,38). NT-dependent and -independent signaling via p75NTR enables this receptor to regulate a broad array of nervous system signaling events (39–41).

A set of pro-apoptotic TNFRs—known as death receptors—possess an ~80 amino acid intracellular death domain (DD) that can engage the extrinsic apoptotic pathway. Homotypic DD interactions with cytosolic death effectors initiate apoptotic signaling cascades that converge on mitochondrial permeabilization and executioner caspase activation (4,42). p75NTR and death receptor 6 (DR6) are atypical DRs that mediate cell death via distinct signaling mechanisms (39,43).

Despite advances to our understanding of TNFR biology, our knowledge of TNFR evolution is limited. TNFRs have been identified in extant species of every animal phylum, but DRs are absent in choanoflagellates and Porifera, suggesting DRs may have originated from a gene fusion event between CRD and DD-containing proteins in an ancestral planulozoan (44,45). *Cf*TNFR—a DR expressed in the Zhikong scallop (*Chlamys farreri*)—shows primary sequence homology to p75NTR in several vertebrate species (46), suggesting that mammalian TNFRs may be descendants of an ancient DR. DRs with sequence homology to p75NTR and EDAR have been discovered in sea anemone, sea urchin, annelids and tunicates (47–49). Moreover, invertebrate chordates express several TNFRs with homology to p75NTR and EDAR, which mediate development of the nervous system and epithelial appendages (44,50,51). Diverse TNFRs with sequence similarity to p75NTR have also been observed in platyhelminthes (52) In *Drosophila melanogaster, a* TNF system that includes one form of TNF (Eiger) and 2 TNFRs [Wengen (Wgn) and Grindelwald (Grnd)] is being increasingly used as a model for TNF/TNFR signaling (53–58). Though individual CRDs of these *Drosophila* TNFRs show homology to several mammalian TNFRs (53,56,58), their evolutionary link to the human TNFR system remains uncertain.

Although TNFRs have been observed in all metazoan phyla (44,45), a comprehensive model of TNFR evolution describing the molecular origins of mammalian TNFRs—and how they relate to the *Drosophila* model system—has remained elusive. To address this problem, we traced the history of all human TNFRs in progressively older extant species spanning the animal kingdom. Our evolutionary tracing focused specifically on the CRD cluster sequence of each TNFR, since this site possesses strong selective pressure with sufficient sequence diversity to distinguish between the receptors. This iterative BLAST strategy provided new insights into the evolutionary history of human TNFRs and identified PITA—a pre-bilaterian TNFR with strong sequence identity to p75NTR—as the common ancestor to mammalian TNFRs. This approach also demonstrated that PITA is conserved all planulozoan phyla and gave rise to the divergent Wgn-Grnd system, thereby establishing an evolutionary link between human TNFRs and Drosophila counterparts.

## Methods

### Evolutionary tracing of human TNFRs

Evolutionary tracing was performed using an iterative BLAST (Basic Local Alignment Search Tool) approach. The CRD cluster sequence for each TNFR was input into BLASTp, queried in the next most ancient extant species, and analyzed according to default algorithm parameters (expect threshold = 10; word size = 6; BLOSUM62 scoring matrix; gap costs – existence: 11 extension: 1; application of a conditional compositional score matrix adjustment; and automatic input parameter adjustment for short sequences). No filtering or masking was applied to the BLASTp query. BLAST hits were identified as *bona fide* homologs if they met the following criteria: (i) E-value ≤ 0.01; (ii) possess 1 or more TNFR CRDs; and (iii) single-pass transmembrane protein. If only 1 protein met these criteria, this protein was identified as a *bona fide* homolog, and the primary sequence that was homologous to the input (within the CRD cluster) was used as the input sequence for the subsequent BLASTp query in the next most ancient extant species. If 2 or more proteins met these criteria, the strongest *bona fide* homolog was identified by meeting these criteria: (i) full-length primary sequence length is most similar to that of the query protein; (ii) BLASTp querying the CRD cluster from other species consistently yields this homolog as the top hit; and (iii) lowest E-value. If no proteins met the basic criteria for homology, the analysis was repeated using the CRD cluster from previous species (as the BLASTp input) in the following order: (i) previous species expressing a homolog; (ii) *Lingula anatina* (if applicable); (iii) a mollusc expressing a homolog (if applicable); (iv) an arthropod expressing a homolog (if applicable); (v) an echinoderm expressing a homolog; (vi) a pre-vertebrate chordate and/or hemichordate expressing a homolog (if applicable); (vii) a teleost expressing a homolog (if applicable); then (viii) the human sequence. If no homolog is identified after these *post hoc* BLASTp queries, we concluded that no TNFR homolog exists in the queried species. If multiple accession numbers are identified for the same homolog, the accession number adhering to standard NCBI nomenclature (e.g. beginning with NP_ or XP_) was selected.

### Identification of PITA, EXT, Wgn and Grnd family members in Protostomia

The CRD cluster sequences PITA and EXT (from *Limulus polyphemus*), Wgn (*Drosophila melanogaster*), and Grnd (*Drosophila melanogaster*) were input into BLASTp and queried in large protostomian taxa, including: Class Insecta, Class Crustacea, Class Chelicerata, Phylum Nematod), and Superphylum Lophotrochozoa (2nd outgroup). BLASTp queries were performed according to default algorithm parameters. Hits with an E-value ≤ 0.001 were identified as bona fide homologs.

### Structural analysis of PITA descendants

Full-length protein sequences for PITA descendants were analyzed for known domains and motifs using the NCBI Conserved Domains Database (CDD) and Pfam (59). Sequence alignments of PITA descandants with human p75NTR were performed in ClustalX version 2.1, and amino acid identity percentages from these alignment were analyzed in Clustal Omega (60). Reconstruction of the known PITA structures was performed manually in Inkscape Version 0.92.

### Evolutionary modelling

All evolutionary modelling was performed using Randomized Axelerated Maximum Likelihood (RAxML) Version 8 software (61). Multiple sequence alignments input into RAxML were generated with ClustalX version 2.1. RAxML output trees were visualized using Interactive Tree of Life (iTOL) version 6.3.2.

### Graphing

Graphs were generated using GraphPad Prism v9.2.0.

## Results

### Human TNFRs are PITA descendants

TNFRs are structurally identifiable by presence of one or more CRDs in their ECD (Table 1). Due the strong selective pressure on these CRD clusters—and the ability to distinguish TNFRs based on primary sequence variation within this region—evolutionary tracing was performed on the CRD cluster of human TNFRs by iterative BLAST queries in extant species with progressively older common ancestry (e.g. the CRD cluster sequence for human TNFR1 was queried in the non-human primate *Pongo abelii*; the homologous CRD region in *P. abelii* TNFR1 homolog was BLAST queried in the non-primate eutherian *Mus musculus*, and so on). Extant species chosen for evolutionary tracing: (i) were selected in an unbiased manner, (ii) span all animal phyla, and (iii) possess a fully sequenced genome (Table 2). As a negative control, the cysteine-rich region (CRR; also known as ‘complement type repeat’) of low-density lipoprotein receptor (LDLR) family members was traced in parallel (S1 Table) due to high sequence similarity to TNFR CRD clusters. The longest tandem cluster of CRRs for a given LDLR family member was selected for evolutionary analysis (S1 Table); importantly, LDLR evolutionary tracing showed no overlap with TNFR evolution.

**Table 1.**
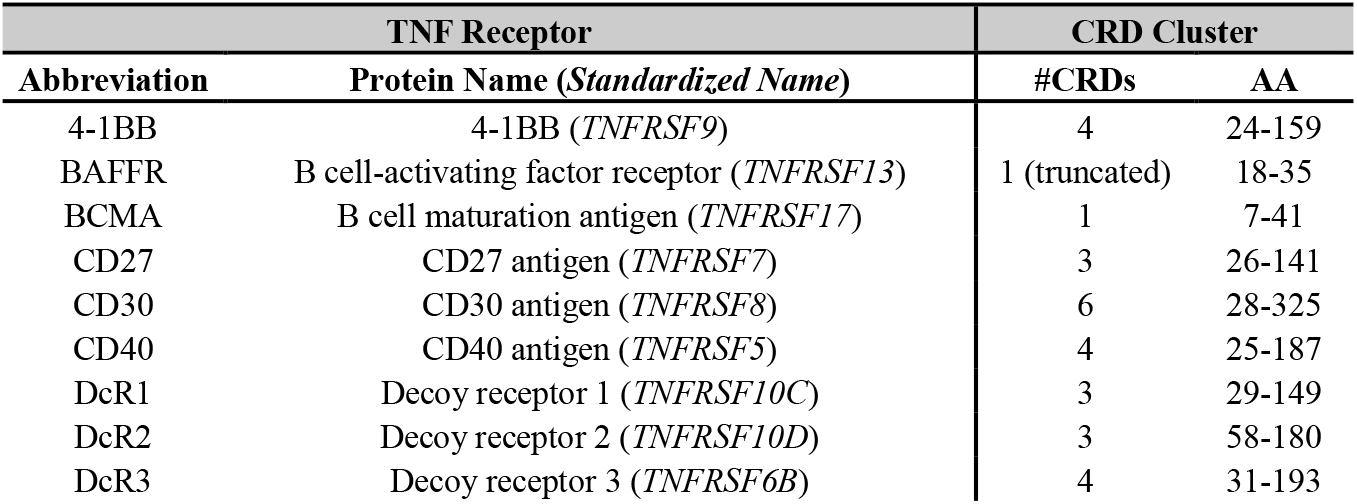
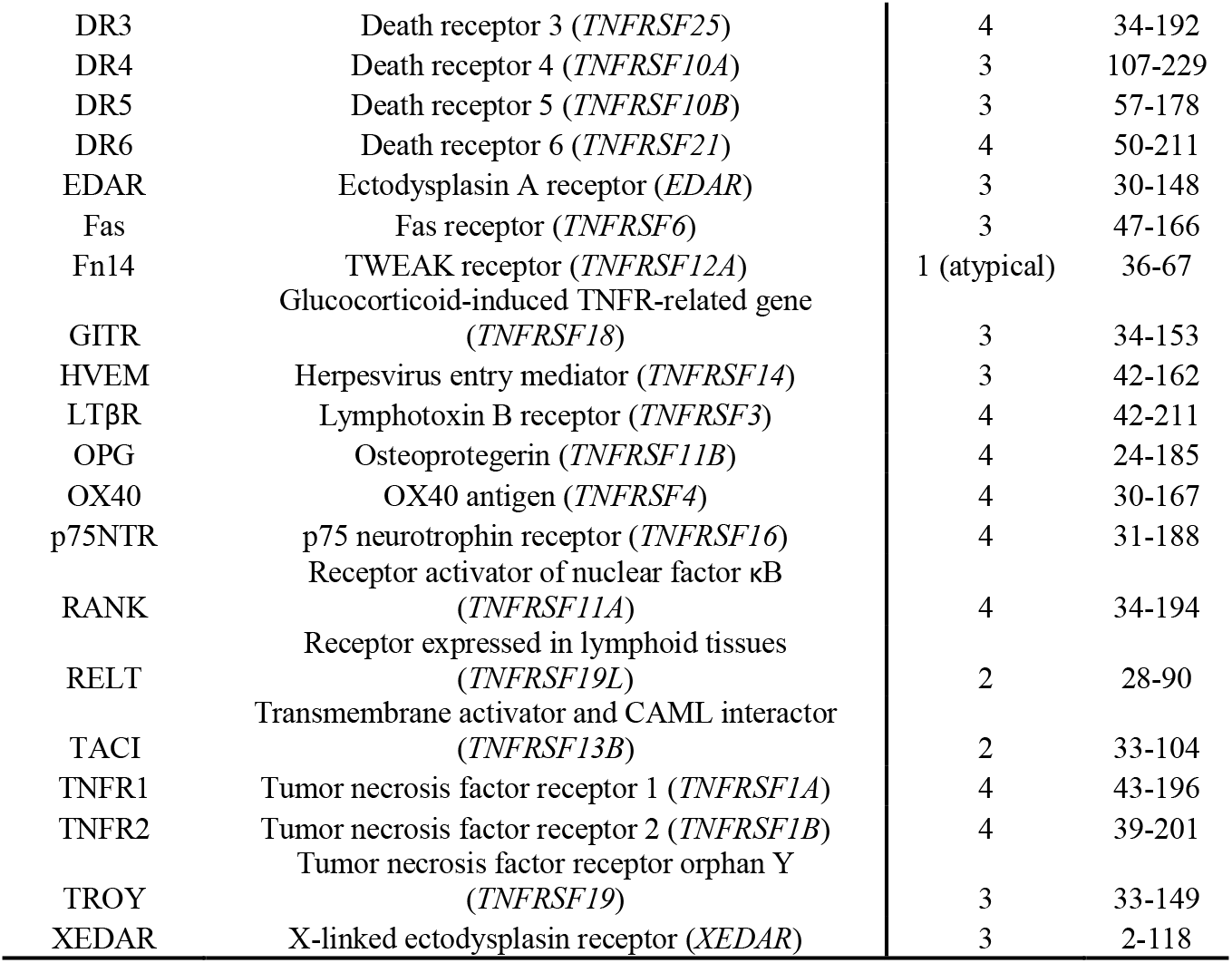
Human TNFRs and their CRD clusters used in evolutionary analyses.

**Table 2.**
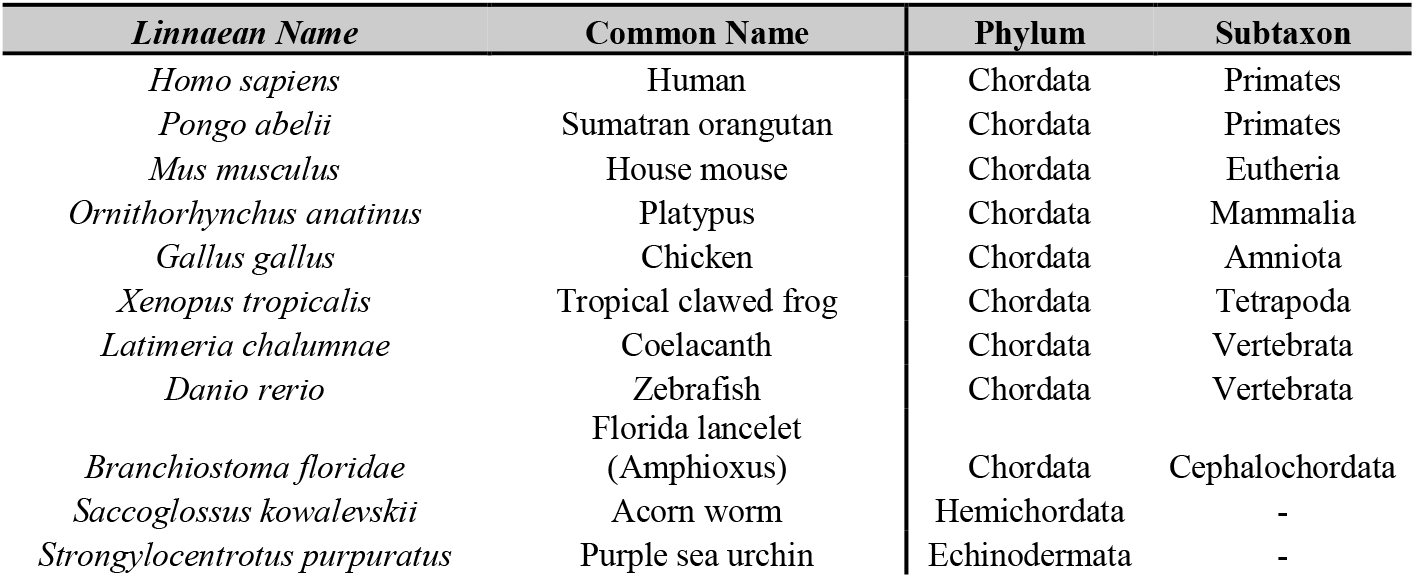
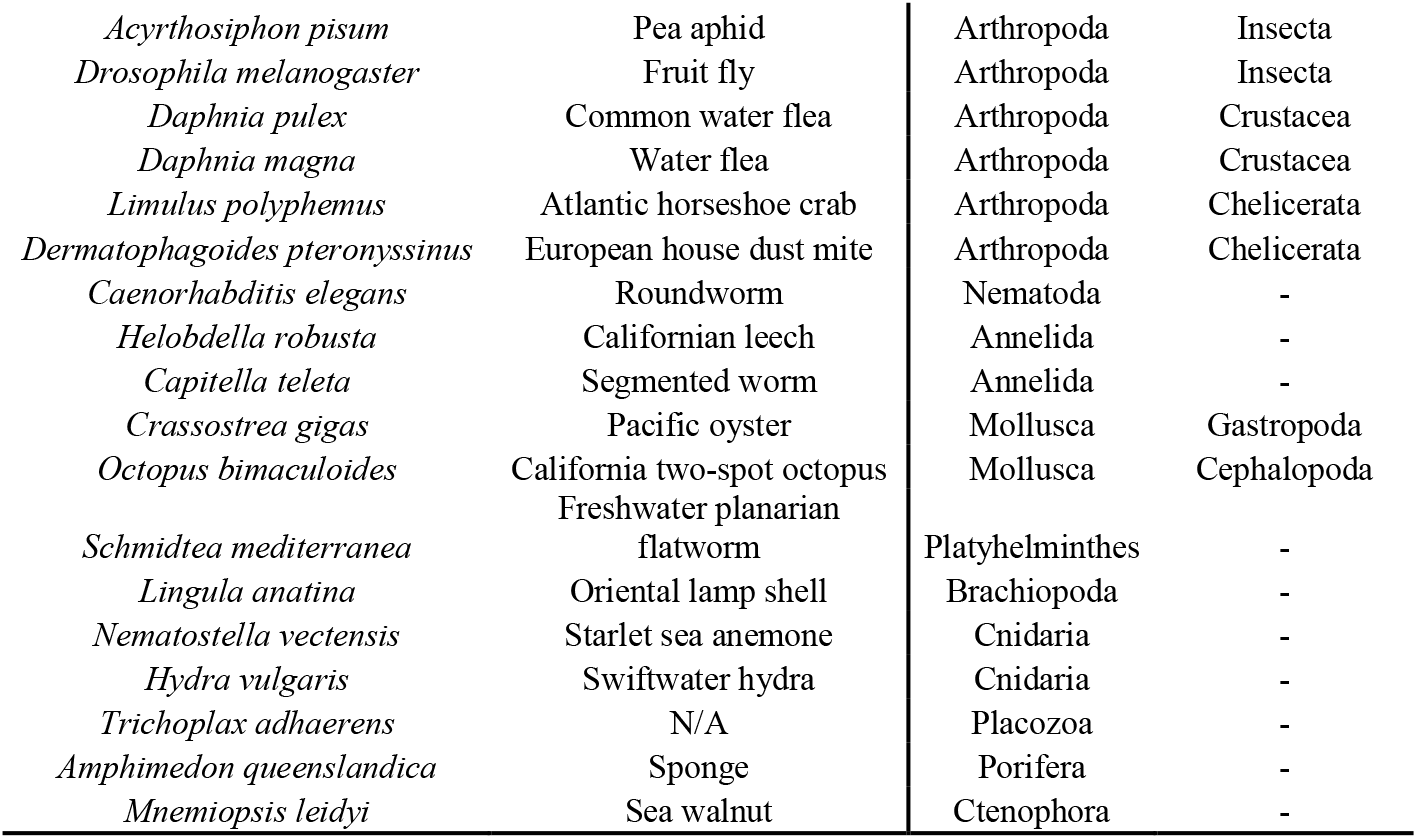
Species used in human TNFR evolutionary analysis.

Remarkably, the ancestry of 25 human TNFRs could be traced to 2 TNFRs expressed in the purple sea urchin (*Strongylocentrotus purpuratus*) (Fig 1A; S1 Table). Of the 25 human TNFRs analyzed, 22 share a common ancestry to a *S. purpuratus* TNFR (XP_030830229.1) that contains 4 CRDs and a death domain. Further tracing of this *S. purpuratus* receptor revealed that orthologs are expressed in a wide array of animal phyla, including Arthropoda, Annelida, Mollusca, Brachiopoda, and Cnidaria (Fig 1A).

**Figure 1.**
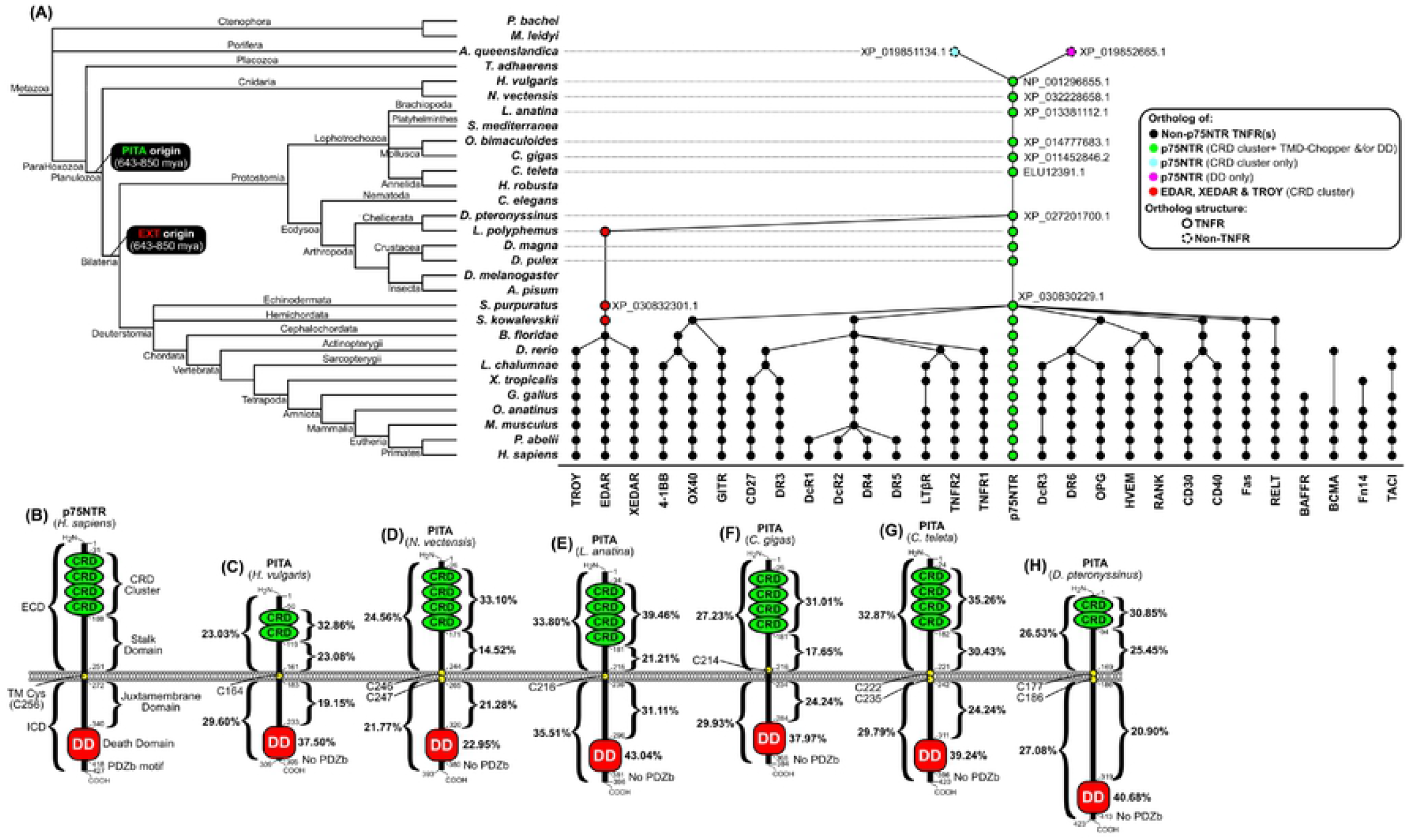
Human TNFRs are PITA descendants. (A) Evolutionary tracing of all 29 human TNFRs. Each dot represents a single homologous protein. A solid outline denotes that the homolog is a TNFR [(i) possesses 1 or more CRDs + a single TMD, and (ii) does not non-CRD functional domains in the ECD]. A dashed outline indicates the protein is not a TNFR [violates criterion (i) and/or (ii)]. The dot’s fill denotes the domains that can recapitulate that homolog (black – a non-p75NTR CRD, green – p75NTR CRD and/or the p75NTR DD and/or the p75NTR TMD-Chopper region, cyan – p75NTR CRD only, pink – p75NTR DD only). (B) Schematic structure of human p75NTR and the domains/motifs/residues analyzed for homology in PITA descendants. (C-H) Structure and % AA identity to human p75NTR (by domain) of PITA descendants expressed in: (C) swiftwater hydra (*Hydra vulgaris*), (D) starlet sea anemone (*Nematostella vectensis*), (E) brachiopod (*Lingula anatina*), (F) Pacific oyster (*Crassostrea gigas*), (G) segmented worm (*Capitella teleta*), and (H) European house dust mite (*Dermatophagoides pteronyssinus*).

The other 3 human TNFRs (EDAR, XEDAR, and TROY) were related to a distinct *S. purpuratus* TNFR (XP_030832301.1) that contains 3 CRDs but lacks a death domain. We identified a single homolog of this *S. purpuratus* TNFR in an arthropod (Atlantic horseshoe crab) but no other orthologs could be identified in lophotrochozoan or pre-bilaterian phyla (Fig 1A). This indicates that the ancestor that gave rise to EDAR, XEDAR, and TROY (designated EXT) originated in an early bilaterian ancestor and has been conserved in protostome and deuterostome evolution. Moreover, this suggests that EDAR independently acquired its DD from a gene fusion event in an ancestral chordate during EXT expansion. Critically, EXT was identified as a descendant of the *S. purpuratus* TNFR that contains 4 CRDs and a death domain (Fig 1A). Thus, all mammalian TNFRs are descendants of a single ancestor conserved in bilaterian and pre-bilaterian phyla.

Four human TNFRs could not be successfully traced to a common ancestor. This is probably because these family members possess 1-2 CRDs that limit the statistical power that applied to their evolutionary tracing. It seems likely that these 4 TNFRs share ancestry to at least 1 of the 2s ancestral TNFRs expressed in *S. purpuratus* — as is the case for the other 25 human TNFRs—but this remains unconfirmed.

We next sought to identify the human TNFR that most closely resembles the sole ancestor identified in Figure 1B. To accomplish this, the full-length sequence of the ancestral TNFR from multiple protostomian and Cnidarian species was BLAST queried in *Homo sapiens*. Reverse BLAST analyses identified p75NTR (10^−9^< E<10^−48^) as the most significant hit (lowest E-value) for every ancestral TNFR sequence queried (Table 3). To corroborate the reverse BLAST result, we ran evolutionary traces on additional p75NTR domains, including the: (i) stalk domain, (ii) transmembrane and chopper domains (TMD-Chopper), and (iii) death domain (DD). Evolutionary tracing of p75NTR by the TMD-Chopper and DD domains replicated the p75NTR ancestry resolved using the CRD cluster analysis (Fig 1A; S1 Table). This demonstrates that p75NTR retained the structural characteristics of the ancestral TNFR. Therefore, we designate the ancestral TNFR PITA, for ***p**75NTR **i**s the **T**NFR **A**ncestor*.

PITA descendants expressed in *Hydra vulgaris* (Fig 1C), *Nematostella vectensis* (Fig 1D), *Lingula anatina* (Fig 1E), *Crassostrea gigas* (Fig 1F), *Capitella teleta* (Fig 1G), and *Dermatophagoides pteronyssinus* (Fig 1H) show strong sequence homology to human p75NTR (Table 3, Fig 1B). Relative to the human p75NTR sequence, PITA sequence conservation is strongest in CRD cluster (30.85% ≤ AA identity ≤ 39.46%) and DD (22.95% ≤ AA identity ≤ 43.04%) and somewhat lower within the intracellular juxtamembrane domain (19.15% ≤ AA identity ≤ 31.11%) and extracellular stalk domain (14.52% ≤ AA identity ≤ 30.43%) (Figs 1C-I; S2 Table).

PITA likely arose from a gene fusion event. To identify extant species carrying domains that contributed to PITA, we traced the p75NTR CRD cluster in pre-ParaHoxozoan phyla using the iterative BLAST strategy. This analysis identified XP_019851134.1 in the Poriferan *Amphimedon queenslandica* (Great Barrier Reef sponge) as a potential origin of the PITA CRD cluster (Fig 1A). Sequence alignment of the CRD clusters from human p75NTR and XP_019851134.1 revealed strong sequence homology (28.32% AA identity) (S3 Fig). A reverse BLAST query of full-length XP_019851134.1 in *Homo sapiens* revealed no human ortholog, but the existence of such an ortholog cannot be ruled out.

Similar evolutionary tracing of the p75NTR DD identified XP_019852665.1 in *A. queenslandica* as a potential origin of the PITA DD (Fig 1A). A reverse BLAST query of full-length XP_019852665.1 in *Homo sapiens* identified this PITA DD ancestor as an ortholog of Death-Associated Protein Kinase 1 (DAPK1) (E<10^−131^). Sequence alignment of human p75NTR DD and the *A. queenslandica* DAPK1 DD revealed significant sequence homology (31.58% amino acid identity) (S4 Fig). Given that XP_019851134.1 and DAPK1 are both expressed in *Amphimedon*, it is reasonable to speculate that fusion of genomic loci encoding the XP_019851134.1 CRD cluster and DAPK1 DD occurred in an ancestral planulozoan. Timetree estimates (62) indicate that this putative fusion event occurred 650-750 million years ago.

Based on the cumulative evidence, we conclude that PITA is the evolutionary ancestor of the human TNFR superfamily. p75NTR retained the ancient PITA structure and all other TNFRs diverged from PITA to acquire unique biological functions. We propose that PITA originated in an ancestral planulozoan from a genomic fusion event between sequences encoding a CRD cluster (XP_019852665.1) and the DD of DAPK1.

### PITAs and EXTs are independent TNFR families

Although all human TNFRs are PITA descendants, EDAR, XEDAR and TROY are descendants of EXT, which diverged in an ancestral bilaterian, earlier than the PITA divergence events in early chordates (Fig 1A). To test if PITAs and EXTs are independent TNFR families, we generated the maximum likelihood phylogeny between PITAs and EXTs (from protostome and deuterostome species) using RAxML. Wgn and Grnd were incorporated into the analysis to clarify the evolutionary relationship between PITAs/EXTs and the Drosophila TNFR model system. To identify PITA and EXT homologs across bilaterian taxa, we performed a large-scale BLAST search using the *L. polyphemus* CRD clusters for PITA and EXT as query sequences (S5 Table). We repeated this strategy to identify Wgn and Grnd homologs by BLAST queries of the *D. melanogaster* CRD cluster sequences with Protostomia. Maximum likelihood modelling of CRD clusters demonstrated that PITAs and EXTs are independent protein families. Importantly, human and murine p75NTR were confirmed as PITA family members, whereas EDAR, XEDAR and TROY were confirmed as EXT family members (Fig. 2A). This indicates that a PITA duplication event in an ancestral bilaterian generated independent PITA and EXT families of TNFRs. In humans, EDAR, XEDAR and TROY are classified as EXT family members; all other TNFRs are classified as PITAs.

**Figure 2.**
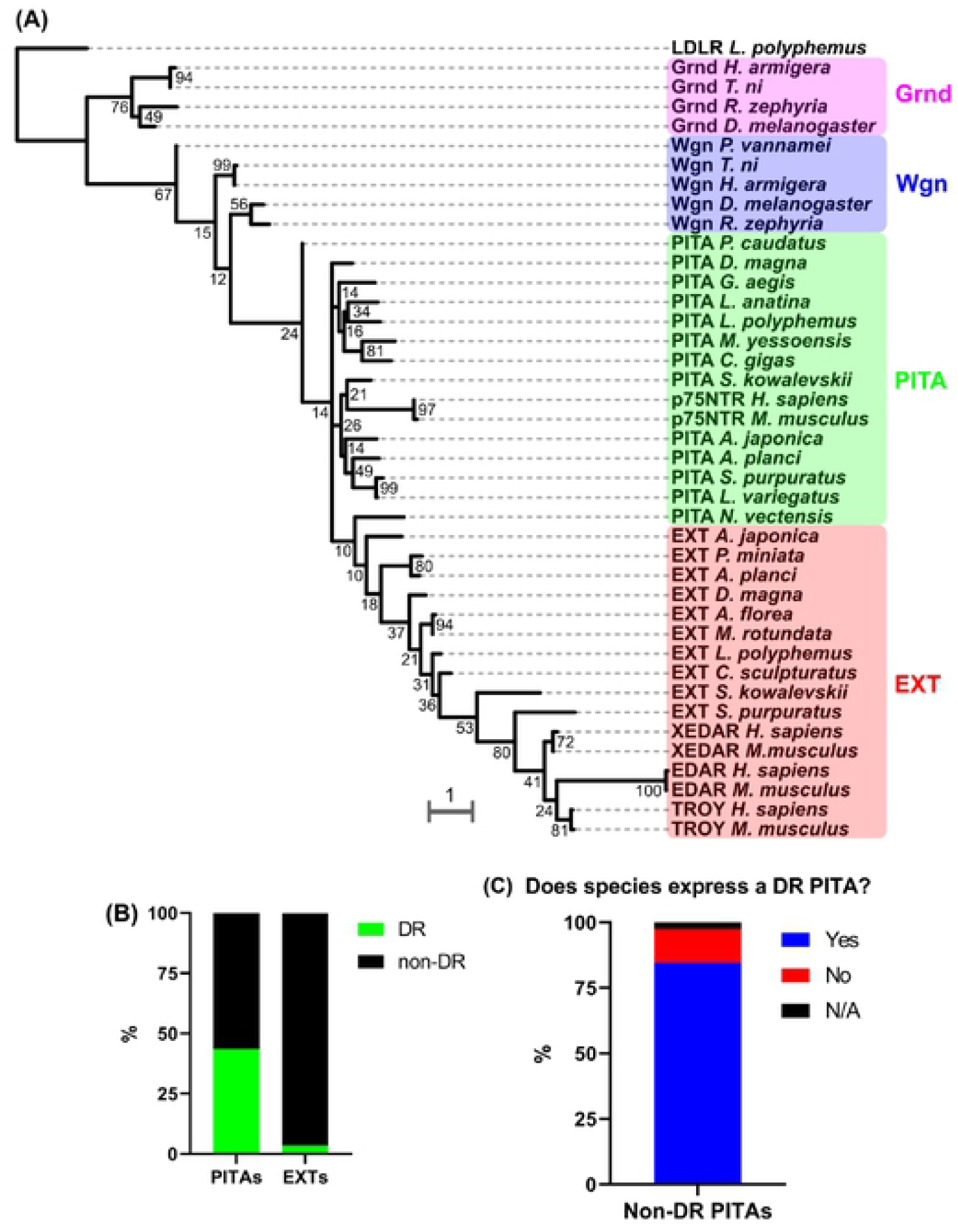
PITAs and EXTs are independent TNFR families. (A) RAxML evolutionary modelling of CRD cluster sequences from PITAs, EXTs, and Wgn/Grnd homologs spanning protostome and deuterostome phyla. The CRR cluster sequence from *L. polyphemus* LDLR was used as an outgroup. (B) Detection of DD in all identified PITA and EXT homologs. Graph represents percentage of protostome PITAs and EXTs where a DD is present (DR) or absent (non-DR). DD search cross-referenced both the NCBI Protein and Pfam databases. (C) Percentage of all non-DR PITAs expressed in species that express at least one DR PITA. ‘Yes’ indicates the non-DR host species expresses a DR PITA; ‘No’ indicates the non-DR host species does not express a DR PITA; and “N/A” indicates the host species’ PITA homologs are only partially sequenced, therefore no determination could be made.

Given that ancestral PITA is a death receptor (Fig 1), we next sought to classify PITAs and EXTs by the presence or absence of a DD. To accomplish this, we BLAST queried the *L. polyphemus* PITA and EXT CRD cluster sequences to identify all protostome homologs. For each identified homolog, we cross-referenced the NCBI Protein and PFAM (63) databases for the presence of a DD (S6 Table). DD-positive homologs were classified as DRs, and DD-negative homologs were identified as non-DRs. Figure 2B shows that 43.5% of all PITAs are DRs, whereas only 3.4% of EXTs possess a DD. To clarify the DR status of the PITA family, we measured the proportion of non-DR PITAs expressed in organisms possessing DR PITA(s). We found that 84.6% of species expressing a non-DR PITA co-express a DR PITA (Fig 2C). These data collectively establish that the vast majority of DRs are PITA descendants, but that rare EXT-descendant DRs also appear in nature (e.g. EDAR in chordates).

### Wengen and Grindelwald are divergent PITA descendants

Wgn and Grnd represent a simple TNFR system assumed to be homologous to mammalian TNFRs. However, the human TNFR evolutionary analysis described in Figure 1 failed to show structural homology of any human TNFR to any protein in the *D. melanogaster* proteome. Given that Wgn/Grnd families are more similar to PITAs than EXTs (Fig 2A), a potential explanation is Wgn and Grnd are divergent PITA descendants that lack direct homology to a human TNFR.

To test this hypothesis, we ran large-scale BLAST queries of CRD clusters from Wgn, Grnd and PITA (*L. polyphemus* sequences) within arthropod subtaxa, including: (i) multiple subtaxa of Class Insecta (Diptera—which includes genus *Drosophila*; Hymenoptera; and ‘All Other Insects’), (ii) Class Crustacea, and (iii) Class Chelicerata. Wgn and PITA homologs were observed in all 3 arthropod classes (Fig 3) and showed no overlap at the single-protein level, with exception of a TNFR expressed in *Amphibalanus amphitrite* (Acorn barnacle) showing weak homology to both Wgn and PITA (E < 0.01) (S7 Table). Grnd homologs were identified within Class Insecta and Class Crustacea, but not Class Chelicerata, indicating that Grnd is an arthropod-specific TNFR originating in a common ancestor to insects and crustaceans (Fig 3). Unexpectedly, a single Wgn homolog was identified outside of the arthropod phylum, in the Nematoda outgroup (CAB3400475.1 in *Caenorhabditis bovis*) (Fig 3; S7 Table), suggesting that Wgn originated in an ancestral ecdysozoa (the clade encompassing arthopod and nematode phyla). (Fig 3; S7 Table). Lophotrochozoa (parallel protostomian clade to ecdysozoa) contain PITAs but do not have Wgn homologs, indicating that Wgn as an ecdysozoan-specific TNFR. A previous study demonstrated that no proteins in *D. melanogaster* nor *Daphnia pulex* —aside from Wgn and Grnd—possess a TNFR CRD consensus sequence (45), ruling out the possibility that an unknown non-PITA TNFR gave rise to the Wgn-Grnd system. This finding, in conjunction with our analyses described above (Fig 2A; Fig 3), indicate that Wgn and Grnd are divergent PITA descendants that share a common evolutionary ancestor with human TNFRs.

**Figure 3.**
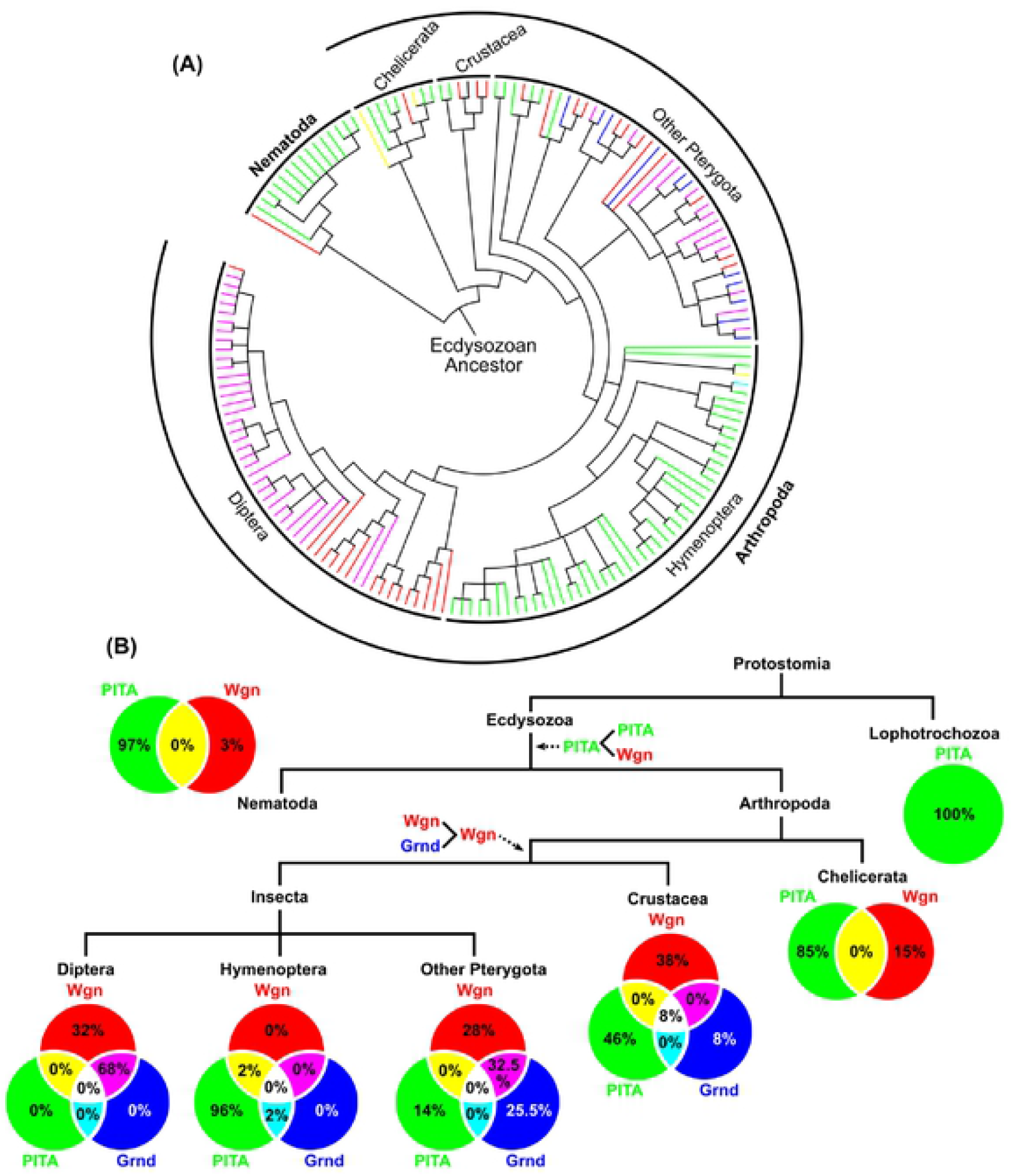
*Drosophila* TNFRs Wengen and Grindelwald are divergent PITA descendants. (A) Phylogenetic reconstruction of all ecdysozoan species expressing Wgn, Grnd and/or PITA homolog(s). BLAST queries were run using the CRD cluster as input (*Drosophila melanogaster* sequences for Wgn and Grnd; *Limulus polyphemus* sequence for PITA). Each species’ TNFR expression profile is represented by the bar colour next to the species name (green – PITA only, red – Wgn only, blue – Grnd only, magenta – Wgn + Grnd, yellow – PITA + Wgn, cyan – PITA + Grnd, black – Wgn + Grnd + PITA). E-value ≤ 0.01 denotes a *bona fide* homolog. (B) Venn diagram summary of the TNFR expression profiles of ecdysoa shown in (A). Lophotrochozoa are included as an outgroup.

Finally, within arthropods, an unexpected trend in the species distribution of PITA and Wgn/Grnd homologs was observed between Diptera and Hymenoptera. Specifically, Dipteran species, including *D. melanogaster*, exclusively express a Wgn/Grnd or Wgn-only TNFR system (no PITAs) whereas Hymenopteran species express a PITA-only TNFR system (no Wgn or Grnd). This suggests that selective pressure may result in mutually exclusive expression of these TNFR systems in Diptera and Hymenoptera.

Based on the available evidence, we propose the following model of Wgn/Grnd evolution: (i) Wgn originated from a PITA gene duplication event in an ecdysoan ancestor; (ii) Grnd originated from a Wgn gene duplication in a common ancestor to insects and crustaceans; and (iii) Wgn and Grnd diverged from PITA to acquire function as receptors to the TNF ligand Eiger.

## Discussion

Using an iterative BLAST approach, we resolved the evolutionary history of the 29 human TNFRs by tracing the origin of the TNFR superfamily to a single ancestral receptor, PITA, that possesses remarkable sequence homology to p75NTR. p75NTR-PITA sequence homology is strongest within the CRD and DD clusters, suggesting that p75NTR and PITA may bind related ligand(s) and activate similar death domain-mediated signaling events. PITA likely originated from a genomic fusion event between the CRD cluster of an uncharacterized TNFR (ancestral to XP_019851134.1 in *Amphimedon queenslandica*) and the DD of DAPK1 in an ancestral planulozoan. According to the Time Tree of Life database (62), the most recent common ancestor between the *Homo* and *Nematostella* genera existed between 647 and 747 million years ago, thus providing an estimate of when this ancestral TNFR originated. PITA is strongly conserved within bilaterian and Cnidarian taxa, indicating it plays a critical physiological role in these species.

Divergence of the PITA lineage occurred first in an ancestral bilaterian, which generated the EXT ancestor; and second, during a massive TNFR expansion event in early chordates. The former event gave rise to an independent TNFR family—known as the EXT family—that proliferated across bilaterian phyla. The latter event coincides with emergence of the adaptive immune system which relies on diverse TNFR signaling events (7). Domain analysis demonstrated that PITAs are commonly death receptors, but EXTs are not, though rare EXT-descendant death receptors do exist. In humans, the closest PITA relative, p75NTR, is a death receptor but most human PITA descendants have lost their intracellular DD. In protostomes, non-DD PITA descendants primarily arise in species already expressing a PITA death receptor. EDAR, XEDAR and TROY are the only EXT descendants expressed in humans. Since non-DR EXTs are prevalent in protostomes, we assume EDAR must have acquired its intracellular DD following an EXT gene duplication event in an early chordate/vertebrate. We suspect the ancestor to human EXTs must have closely resembled XEDAR or TROY, but this is not definitive; neither receptor shows strong conservation to the ancestral sequence, as is observed with p75NTR and ancestral PITA.

Clues to the functional origin(s) of the ancestral TNFR may come from the observation that PITA descendants first appear in the phylum Cnidaria which includes corals, jellyfish and sea anemones. Morphological features appearing in Cnidaria include tissue layering, radial symmetry, and synaptic neural networks (64). Importantly, the placozoan *Trichoplax adhaerens*—which lacks adaptations that lead to fast synaptic transmission (e.g. synaptic cleft, synaptic vesicles, etc.) (65)—does not express a PITA descendant. Thus, PITA expression may be limited to animals possessing synaptic neural networks. This in turn suggests that the emergence of PITA may have been a prerequisite for the development of complex nervous system networks. Future investigation into the biological function of PITA in a Cnidarian model organism, such as the sea anemone *Nematostella vectensis*, may yield insight into the core functionality of this TNFR ancestor.

## Conclusions

This study describes the first comprehensive model of human TNFR evolution and identified the death receptor PITA—a novel TNFR with strong sequence homology to p75NTR—as the molecular ancestor to the mammalian TNFR superfamily. PITA was likely created by the fusion of genomic loci encoding the CRD cluster of the uncharacterized TNFR XP_030830229.1 and the DD of DAPK1 in a common ancestor to humans and the demosponge *Amphimedon queenslandica*. The PITA receptor has remained structurally intact over 750 million years of evolution and gave rise to several descendent TNFR families in deuterostomes and protostomes, including the *Drosophila melanogaster* TNFRs Wgn and Grnd.

## Supplemental Information

**S1 Table. Raw TNFR and LDLR evolution data.** The NCBI accession number and homologous domain sequence for each identified TNFR/LDLR homolog.

**S2 Table. Sequence identity of PITA descendants to human p75NTR by protein region.**

**S3 Figure. Structure of XP_019851134.1 in *Amphimedon queenslandica*.** (A) Structural schematic of *A. queenslandica* protein XP_019851134.1 (shown to scale). (B) Multiple sequence alignment of the CRD cluster sequences for XP_019851134.1 and human p75NTR.

**S4 Figure. Structure of *Amphimedon queenslandica* DAPK1 ortholog.** (A) Structural schematic of *A. queenslandica* DAPK1 (shown to scale). (B) Multiple sequence alignment of the death domain sequences for *A. queenslandica* DAPK1 and human p75NTR.

**S5 Table. Large-Scale BLAST queries of PITA and EXT in ecdysozoa and lophotrophozoa.**

**S6 Table. PITAs, EXTs, and Wgn/Grnd homologs spanning bilaterian phyla used for evolutionary modelling.**

**S7 Table. Large-scale BLAST queries of Wgn, Grnd and PITA in protostomian taxa.**

**S8 Figure. Full-length sequence alignments of PITA descendants and human p75NTR.**

